# *crossword*: A data-driven simulation language for the design of genetic-mapping experiments and breeding strategies

**DOI:** 10.1101/330563

**Authors:** Korani Walid, Justin N Vaughn

## Abstract

The simulation of genetic systems can save time and resources by optimizing the logistics of an experiment. Current tools are difficult to use by those unfamiliar with programming, and these tools rarely address the actual genetic structure of the population under study. Here, we introduce *crossword*, which utilizes the widely available results of re-sequencing and genomics data to create more realistic simulations and to simplify user input. The software was written in R, making installation and implementation straightforward. Because crossword is a domain-specific language, it allows complex and unique simulations to be performed, but the language is supported by a graphical interface that guides users through functions and options. We first show *crossword*’s utility in QTL-seq design, where its output accurately reflects empirical data. By introducing the concept of levels to reflect family relatedness, *crossword* is suitable to a broad range of breeding programs and crops. Using levels, we further illustrate *crossword*’s capabilities by examining the effect of family size and number of selfing generations on phenotyping accuracy and genomic selection. Additionally, we explore the ramifications of effect polarity among parents in a mapping cross, a scenario that is common in crop genetics but often difficult to simulate. Given the ease of use and apparent realism, we anticipate *crossword* will quickly become a “bicycle for the [geneticist’s] mind”.

## Introduction

The simulation of controlled crosses has been useful in both applied breeding (reviewed in [1]) and genetic mapping [2,3]. Current open-source packages have expanded the realism and utility of simulated scenerios by incorporating elements such as variation in recombination frequencies, novel transgenic approaches, and genomic selection, among others [4,5]. Still, this realism comes at a significant cost, both, in terms of computational speed and difficulty of use. Using emerging genomics data, we have developed a simulation platform that combines ease-of-use with highly realistic behavior. The platform, called “*crossword*”, is essentially a domain-specific language that, when executed, is interpreted into and executed by the R statistic programming environment. This layer of abstraction allows users to focus *less* on the mechanics of implementing the simulation and *more* on their actual experimental goals and/or breeding ideas. In addition, this precise syntax allows *crossword* to take full advantage of the familial structure unique to controlled crosses.

While the capabilities of simulators such as AlphaSim are substantial, it and other similar packages still do not take full advantage of empirical genomics data that could substantially simplify both the computation and usage of simulation for plant and animal breeders. For example, the first step in most simulation frameworks usually involves generating a large-number of founder haplotypes via coalescent simulation with recombination. Parents in the pedigree are then randomly selected from these founders. In the very near future, breeding programs will also have access to genome sequences for many founder lines within a program as well as extensive genotypic information to impute this information onto their progeny populations. Thus, initial coalescent simulations that create founder sequences are often irrelevant and even an obstacle for many breeders, who already have resequencing data for the parent lines in their study and would prefer to use that information. These unrealistic simplifying assumptions about founder structure can also have substantial implications on predicting breeding results [6]. Although, some of these simulators offer a means to supply parental genotypes, this functionally is very difficult to implement in practice. Only recently have some researchers begun to build in functionality for direct import of genotypic data [6]. Indeed, to our knowledge, no simulation framework fully supports the integration of empirical data, such as genome sequences and annotations, that are currently available for most crop species. *crossword* has native support for importing FASTA, GFF, VCF, and HAMAP file formats and uses this information to substantially simplify the user experience and make simulations as case-specific as possible.

*crossword* has three anticipated tiers of usage: 1) Most users will initially interact with *crossword* via a semi-graphical editor run under gWidgets/tcltk, a lightweight R windowing environment (Figure 1A). This interface frees users of having to thoroughly familiarize themselves with the functions and parameters prior to use. Commands are automatically generated based on either the displayed defaults or the values supplied to the text-box entries. 2) All *crossword* runs generate a plain-text, run script. Therefore, those who wish to automate the exploration of a large-range of parameters can manipulate a previously-generated template by manually replacing parameters of interest with variable names and supplying a range file when running the modified script. 3) All R functions underlying *crossword* are available and well-documented for those who have experience in R and need to directly modify or expand a behavior. Often, we find that many users interact with *crossword* in all three of the above ways.

**Figure 1:**
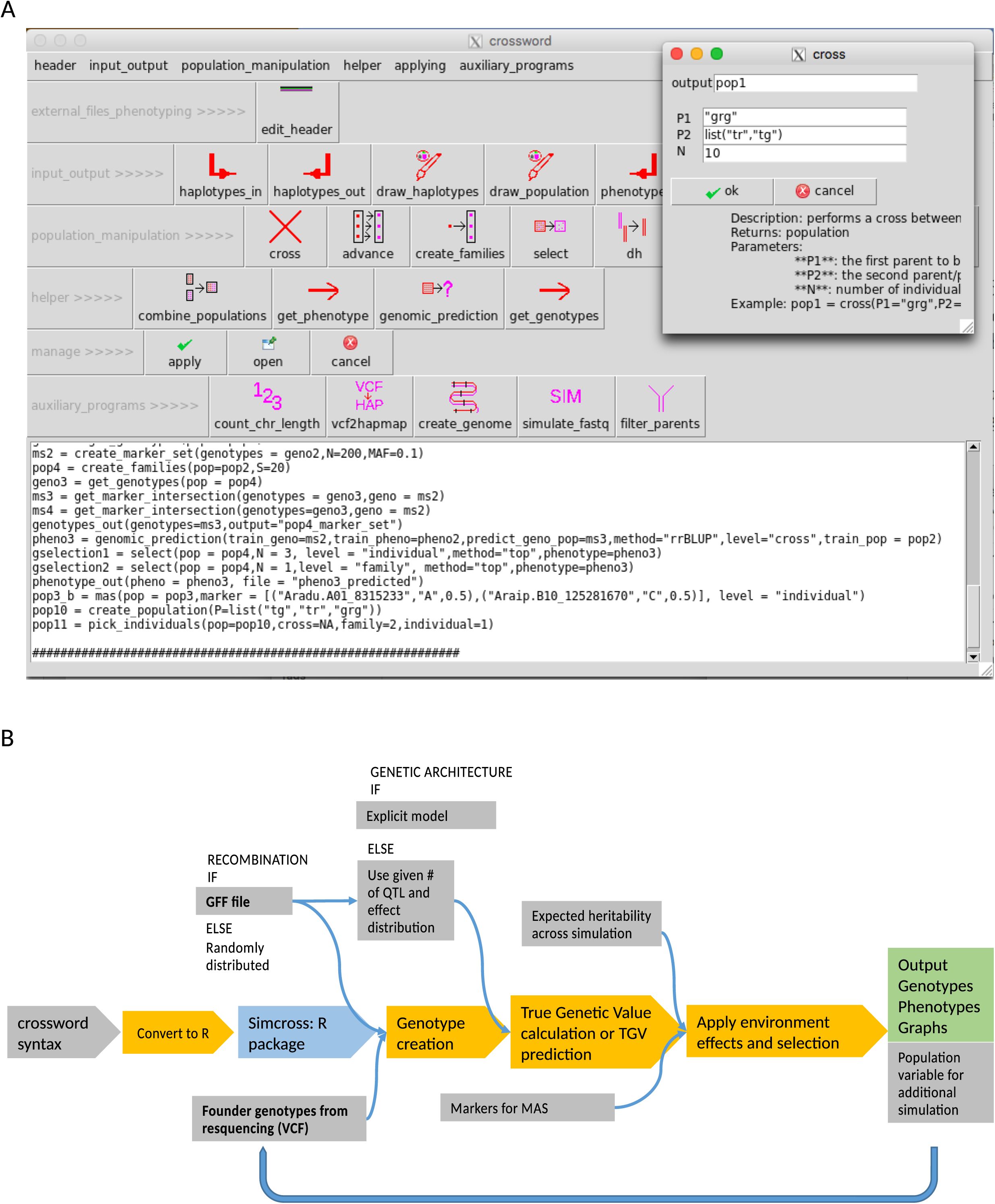
Syntax and execution of a crossword run. A) User interface layout and editing window for crossword syntax. B) Schematic of crossword simulation execution. Gray color indicates arguments or files taken from user. Blue fill indicates an external dependency. Bold font indicates major source of external genomic data.

## Methods

### Genetic Architecture

The genetic architecture of traits is simulated in one of three modes: *QTN_random, QTN_supplied*, or *high_low_parent*. In *QTN_random* mode, each QTN position is drawn at random across the genome and underlying allele effects are assigned based on the supplied distribution. In *QTN_supplied* mode, positions and the effects associated with alleles are defined by the user. In *high_low_parent* mode, postions are drawn at random, as in *QTN_random* above, but allelic effects are biased to be positive in the user-specified high parent and negative in the low parent. True genetic values (TGVs) are then summed for a particular genotype. (If QTN are dominant, the TGV and true breeding value will be different.) Epistatic interactions are currently unsupported.

In its simplest form, heritability (H^2^) is the relationship between additive genetic variance (V_G_) and residual variance (V_R_). V_G_ can change substantially depending on the parents in a cross and downstream selection in a population, among other things. Alternatively, V_R_ will be essentially constant assuming individuals are evaluated in a consistent set of environments. The common approach of calculating V_R_ from the supplied H^2^ for every phenotyping step is therefore conceptually dubious, since we do not expect V_G_ and V_R_ to show substantial covariance. Indeed, in long-term selection experiments, the conventional approach will dramatically overestimate the ability to accurately select superior genotypes in late generations. To that end, in *crossword*, the user supplies either an “absolute” H^2^, defined as the heritability in a theoretical cross that segregates for all possible QTN, or an “average” H^2^ across all possible crosses between all founders. (If there are only two founders, then the models are equivalent.) In these theoretical progeny populations, we assume a very large population size, free recombination, and complete homozygosity. V_R_ is then calculated using an approximation function created from 10,000 individuals binomially distributed for positive and negative alleles. The approximation is used to simplify calculations when effects are not equal and/or do not follow a formal distribution. The actual H^2^ in any one population is reported whenever that population is phenotyped, such that the user can modify the supplied H^2^ if it conflicts with their expectation.

### Founder haplotypes and recombination

In heterozygous founders, we assume that the haplotype phase is unknown and randomly select one allele at heterozygous SNPs to belong to one homolog. Heterozygous sites in founders are discarded when homo=TRUE, which is default behavior because heterozygosity in highly inbred founders is often a genotyping error. When homo=FALSE, heterozygous loci are randomly phased into parental haplotypes. Recombination between founder haplotypes is simulated using the external R library, simcross, which takes into account interference (m=10, by default). By default, at least 1 chiasma per meiosis is required. Supplemental Figure 2 shows the genomic loci of crossing-over across different population size.

**Figure 2:**
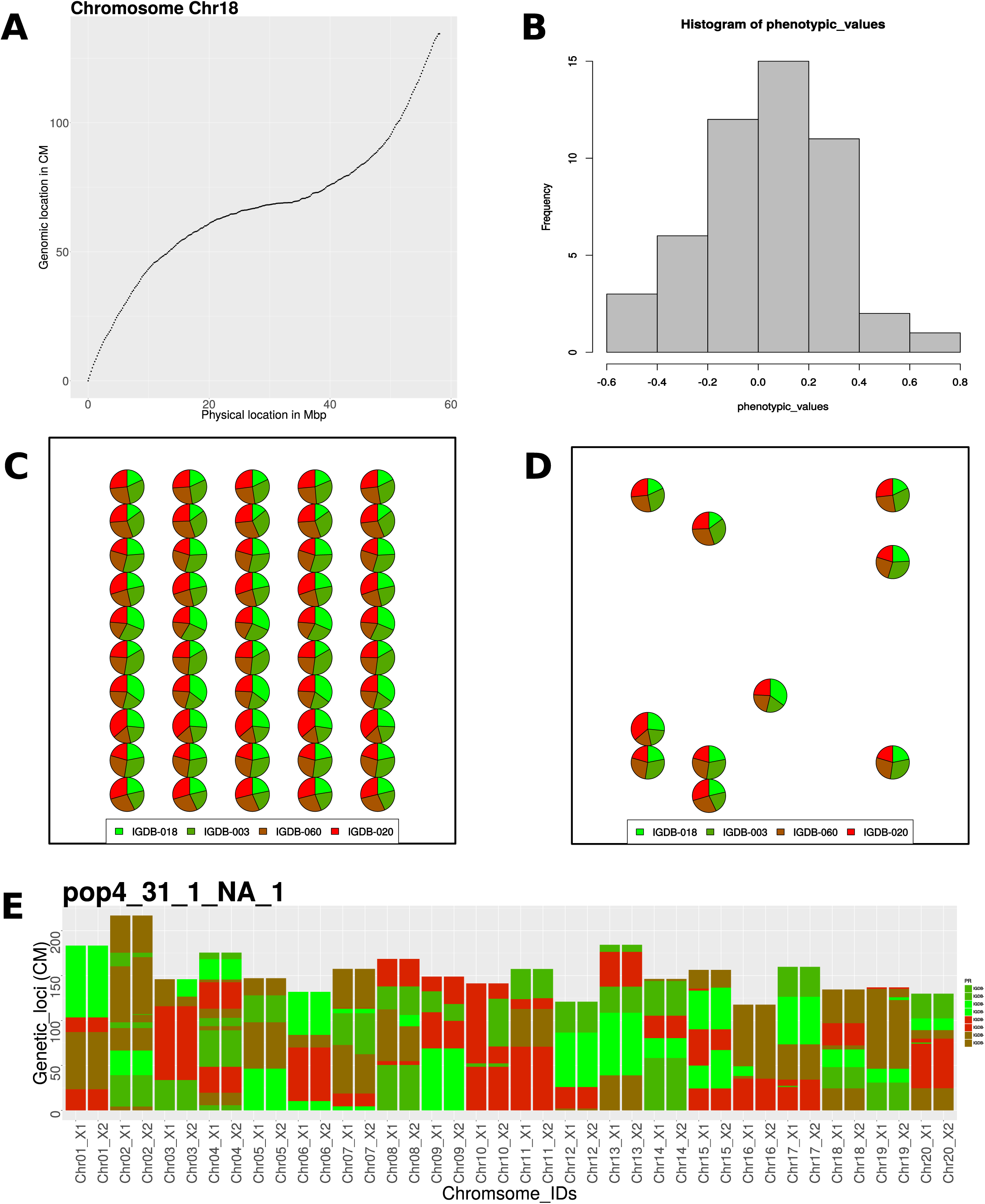
Graphical output menagerie. A) Relationship of physical distance (Mb) to genetic distance (cM) based on gene density in a sliding widows of 10 Kbp. B) Phenotypic distribution of a particular population. C) Parental contributions of the population resulting from 4 founders intercrossed and advanced for F5 generations. The population is composed 10 families, each having 5 individuals. D) Chromosomal representations of an individual from one of the above families. X1 and X2 represent the homologous chromosomes. Identical-by-descent regions are color coded based on founding donors.

Founder haplotypes are imported in either HapMap or VCF formats. The names of founders in these files can be used directly in *crossword* to define parents. As with most contemporary simulations [7], only haplotype structure is simulated under a given set of crossing/advancement steps. When phenotyping or genotyping is performed, physical-to-genetic distance functions are used to produce genotypic information for each haplotype. The relationship between genetic and physical distances is known to be nonlinear for the vast majority of species [8]. In *crossword*, we use the known relationship between gene density and crossover frequency to produce a function that can be used as a substitute for explicit mapping studies. GFF files are supplied to *crossword*. A sliding window is used to calculate gene density versus physical distance. The the R “loess” function is fit to the resultant relationship and this function is used to convert physical coordinates into genetic coordinates based on total genetic length (cM) supplied by user or 100cM for all chromosomes by default.

### Genetic variance in founder population versus single cross

All polymorphisms in the input founder file (VCF or hapmap) are used to select QTN. Therefore, when making a cross, the actual number of segregating QTN will be lower if the founder file contains more than the two individuals in the cross. This approach allows users to apply genome-wide association data from the diversity panels that they may be using as founders and to accurately simulate the reduction in relative genetic variance that occurs in derived populations. A helper function is supplied to assist in reducing VCF files depending on the level of polymorphism desired.

### Genomic prediction

The genomic estimated breeding values (GEBVs) for genotypes are predicted using rrBLUP and the supplied training data. Currently, only rrBLUP is supported, but, because *crossword* is based on user-supplied genotyping files, discontinuous runs can be easily implemented by exporting genotypic information at one stage and importing in a separate round of simulation. This gives users the flexibility to use or test their own tools without having to rely solely on those that we have directly integrated. (Users with experience in R can also directly modify or augment the underlying algorithms to avoid discontinuous simulation.)

### QTL identification and effect estimates for biparental cross

In simulations, QTL where mapped using r/qtl [9]. To focus on effect estimates, the true genetic positions for all markers were used directly as opposed to being estimated from recombinants. Using ‘stepwiseqtl’ function, ‘max.qtl’ were iteratively increased until the minimum LOD score across all discovered QTL was <3. Effects were assessed based on the final resultant model. The R script to perform mapping is available in the auxillary_scripts folder released with *crossword*, but depends on r/qtl, which is not included with *crossword*.

### QTL-seq genetic resolution determination

Data and parameters are derived from [10] and (Joshua Clevenger, personal communication). The genomic data of peanut and soybean was collected from PeanutBase (https://peanutbase.org) and Phytozome (https://phytozome.jgi.doe.gov), respectively. The genetic map information was collect from David Bertioli and Ben Stewart-Brown (personal communications), respectively.

*crossword* offers the capability of simulating reads from a genomic sequence generated from a set of individuals in a simulation. Reads can then be simulated from these sequences if a user wanted to explore the full pipeline of a genotyping-by-sequencing pipeline. Alternatively, since this read simulation approach can be computationally demanding, *crossword* offers an auxiliary program to add sampling and read coverage variation to a set of underlying genotypes. In actual experiments involving bulking, such as QTL-seq, these genotypes are not known, but in a simulations users can exploit this knowledge to get a realistic sampling of each bulked sample. In our simulation, we used the average coverage from the original experiment (30x across the bulk) and a standard deviation of 7. For each locus, a random variant was sampled from a normal distribution based on these values, and this random number of alleles was sampled with replacement from the set of all bulked genotypes.

### Availability

*crossword* is available at https://github.com/USDA-ARS-GBRU/crossword. A detailed manual, function description file, links of youtube tutorial videos and demo data are available at the same link. Each simulation described in this manuscript is also available as an example (see https://github.com/USDA-ARS-GBRU/crossword/examples) identified by the figure to which it applies. Users can submit requests for functionality at.

## Results and Discussion

### Graphical interface and output facilitate “out-of-the-box” useability and utility

The *crossword* run script is interpreted into a set of R functions; thus, *crossword* is easy to install on any system capable of running R. *crossword* allows the user to submit a text file defining a small number of input files and describing the crossing scheme in a simple but powerful syntax (Figure 1A and Supplemental Figure 1). A semi-graphical interface guides users to the appropriate functions, reduces typing errors, and supplies “point-of-use” documentation.

The founder haplotypes in any *crossword* simulation are supplied by the user as either a VCF or HapMap file (Figure 1B). Large population sizes coupled with numerous breeding cycles can be computationally burdensome if using genome-wide polymorphism data. Instead, as with most current simulators, *crossword* only tracks recombinations and the relevant haplotypes [7]. During simulation the entire founder file in scanned for polymorphisms. For example, if the genetic architecture consists of 30 QTN, then these QTN are sampled from across population-wide polymorphisms and so a single cross may or may not be segregating for all 30 QTN. If a user wants all specified QTNs to be segregating in a biparental cross, they can filter VCF files to the appropriate parents. (*crossword* supplies auxiliary scripts to perform this and other capabilities, if needed). Only when a phenotyping or genotyping step is performed on a population are genetic coordinates converted to genomic coordinates (see below) and phenotypes calculated. Given the computational demand on this step, *crossword* uses the Rcpp external R library and custom C++ programs the speed operations ~60-fold.

Users are able to generate graphical output depicting haplotypes for any individual at any generation, genetic-by-physical distance functions for each chromosome, genetic breeding values relative to phenotypic simulation, and dense plots of population structure and founder contributions for a population (Figure 2). Numerous output formats are supported including SVG and PNG.

### Gene-density approximation accurately reflects generic crossover frequency

One of *crossword*’s most useful characteristics relates to its usage of pre-existing genomic information to increase the realism of simulations without overburdening the user. As an example, other simulation suites, if they allow variation in recombination along the genome, require the user to manually create genetic-by-physical distance matrixes in order to simulate this behavior. This requires access to data from a particular mapping study or substantial effort to align markers to the genome. As an alternative, *crossword* takes advantage of the broadly observed correlation between gene density and recombination rate [8] in order to create this function. Gene density can be easily acquired from a GFF file available with almost any genome release. Thus, the user merely has to download said file and define its location in *crossword*.

We compared the performance with two genetic maps, one in peanut and one in soybean (David Bertioli and Ben Stewart-Brown, personal communication). Given that these represent only two populations within each species, the fit generally approximates the maps with the exception that the centromere displays slightly less recombination than predicated by gene density alone (Figure 3). Still, we feel the tradeoff for ease-of-use justifies this minor discrepancy.

**Figure 3:**
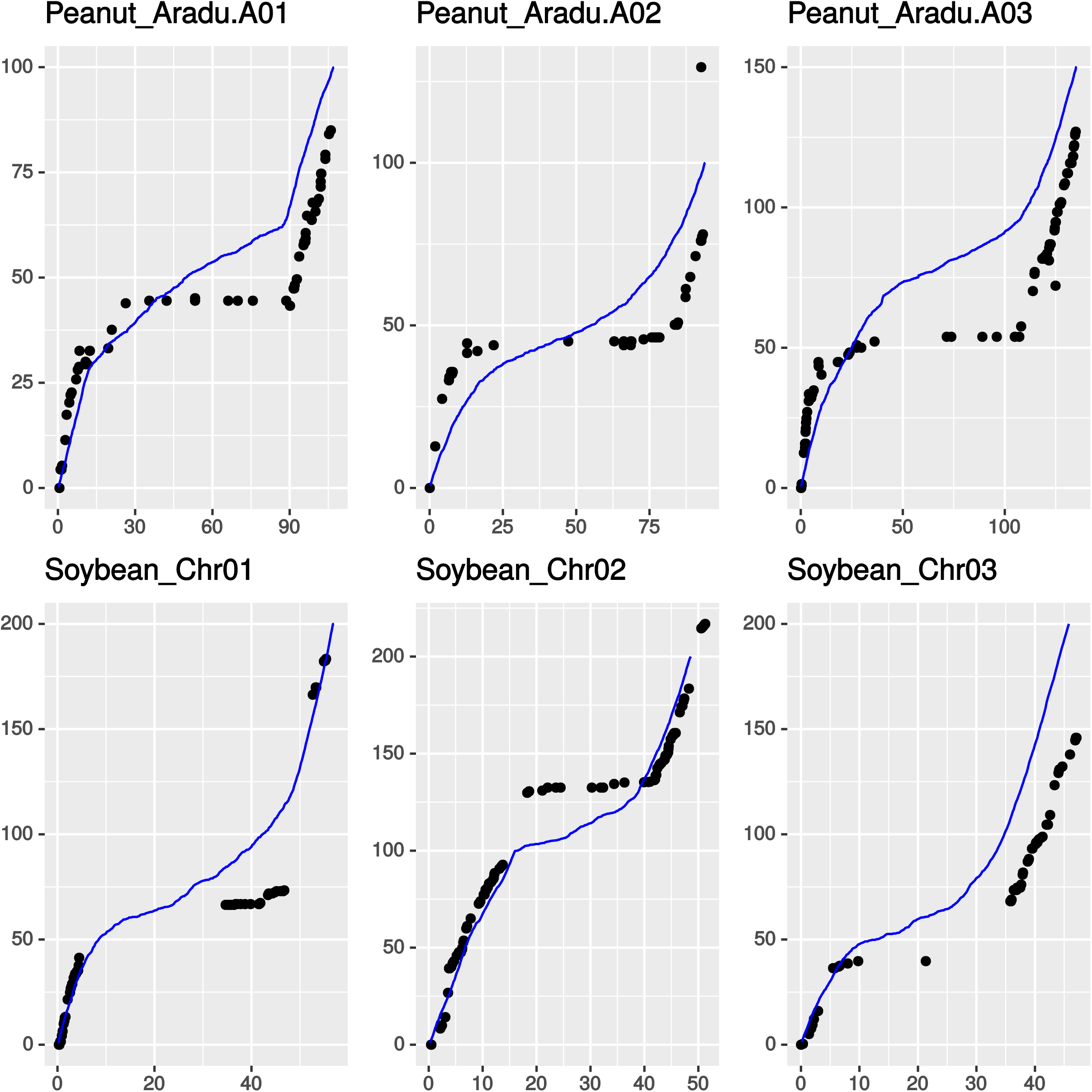
Recombination modeling in crossword. The relationship between genetic and physical distance is shown for three randomly selected chromosomes from two species. Points are based on empirical data from biparental populations whereas lines represent the function used by crossword base on gene density.

In addition, gene density is almost certainly correlated with causal variants (or quantitative trait nucleotide (QTN)) density for particular traits. Instead of randomly picking positions across the genome, *crossword* can use this gene density information to bias randomization to fit this more realistic model.

### Empirical and crossword results for QTL-seq on oligogenic disease resistant trait

Genetic mapping is a central goal of many research programs. With the advent of CRISPR/Cas9 methodologies and the availability of extensive resequencing data, the emphasis in many experiments has shifted from rough QTL identification to causal gene or functional marker discovery. Regardless of the objective, any expectation of genetic resolution is beneficial to experimental design. While “back-of-the-envelope” calculations can be done in terms of expected recombinations given a certain population size, subtler details, such as phenotyping accuracy, residual heterozygosity, and unequal effect sizes can substantially affect intuitive estimates. These complications are exacerbated with increasing number of QTN.

QTL-seq is an emerging genetic mapping strategy that combines bulk segregent analysis with next-generation sequencing. We simulated one such previously published QTL-seq experiment and included all parameters relevant to that population and its development ([10] and Joshua Clevenger, personal communication).

The empirical data revealed two strong peaks at chromosomes Aradu.A05 and Araip.B05, and one weak peak at chromosome Araip.B03 correlated with late leaf spot resistance of peanut. The H^2^ could not be determined from QTL-seq data, so we simulated both h2 = 0.5 and 0.8. The study used peanut genotypes SPT06-06 as a resistant Florida07 as a susceptible with 5514 markers segregating. A population of 166 individuals was advanced to F6. The highest and lowest 5% was bulked for QTL-seq analysis. A simulation was run using the same parameters, which showed similar trends for the three peaks (Figure 4A). However, using lower H^2^ showed results more closely resembling the empirical data (Figure 4B). In addition, we used equal, while the empirical data indicate at least one QTL (on B03) is quite minor.

**Figure 4:**
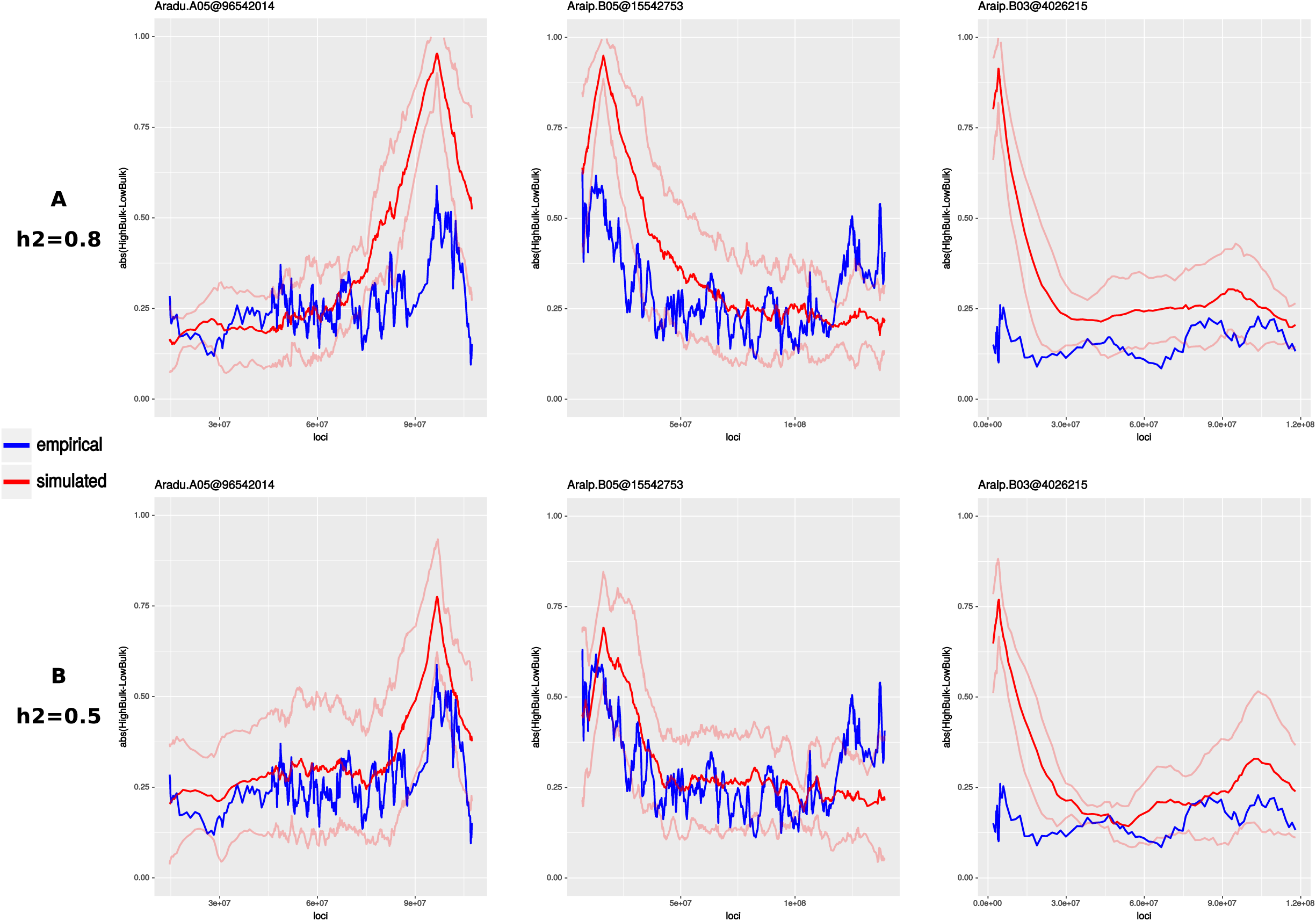
Empirical versus simulated QTL-seq results, red lines show the genetic resolution for QTL based on 10 iterations in which parameters reflect those of the actual experiment, except that all effects were simulated as equal. Blue lines indicate the genetic resolution of the empirical results. For all sharp lines represent the means and faded lines represent the standard deviation. A) H^2^ = 0.8 B) H^2^= 0.5.

### “Levels” in crossword: genotypic replication in phenotype estimation and genomic selection

A major aim of *crossword* is to supply a flexible but powerful syntax for the exploration of realistic breeding strategies. Previous simulation frameworks have underestimated the importance of how particular genetic pools get advanced and evaluated and, therefore, underestimating the impact of factors such as residual heterozygosity and family size on genetic variance and phenotypic replication. For example, bulk advancement versus single-seed descent can have a substantial impact on the logistics of line management. How much impact will the resultant genetics have on downstream breeding targets? Does the cost of reproductively isolating lines outweigh the impact of low outcrossing rate? Some population level processes can be modeled using scripting libraries such as simuPop [11], but scripting requires a general proficiency in programming and, more importantly, is not designed for dealing directly with controlled crosses. We have addressed this operational gap by incorporating the concept of “levels” into many *crossword* functions. Where applicable, users can specify one of four levels: “individual”, “family”, “cross”, or “population”. All populations have “individual” and “population” levels, but other levels depend on how the population is created and managed (See Supplemental Figure 1). For example, a researcher may want to create a population by crossing one individual by three other lines and then advancing the entire population in bulk. Alternatively, they may want to advance each cross as a separate bulk. The level parameter in *crossword* allows these variations with very minor changes when calling the “advance” function.

The “level” concept carries over into phenotyping and selection. Generally, the costliest aspect in genetic mapping and breeding is phenotyping. Unless clones or double haploids are used (which *crossword* supports), nearly all phenotypic replication of a “genotype” is actually based on a population of closely related individuals, usually referred to as a “recombinant inbred family”. *crossword* gives users the power to fine-tune this family-creation function in order to precisely explore the implications of sample size on their final result.

To further illustrate the level concept, we explored the trade-off between time, field space/resources, and phenotypic accuracy. We simulated multiple combinations of generations and family sizes (Figure 5A). For each line we compared the estimated true genetic (TGV) value calculated from family replicates to the TGV of the family progenitor. As expected, the accuracy of TGV improves as a result of both family homogenization - generations of selfing - and family size. Still, the degree to which these factors interact and the optimal balance between accuracy and expense naturally depends on the constraints of the research/breeding program. Clearly, there are diminishing returns of phenotypic accuracy as the number of replications goes up, but this diminishment is a function of generations.

**Figure 5:**
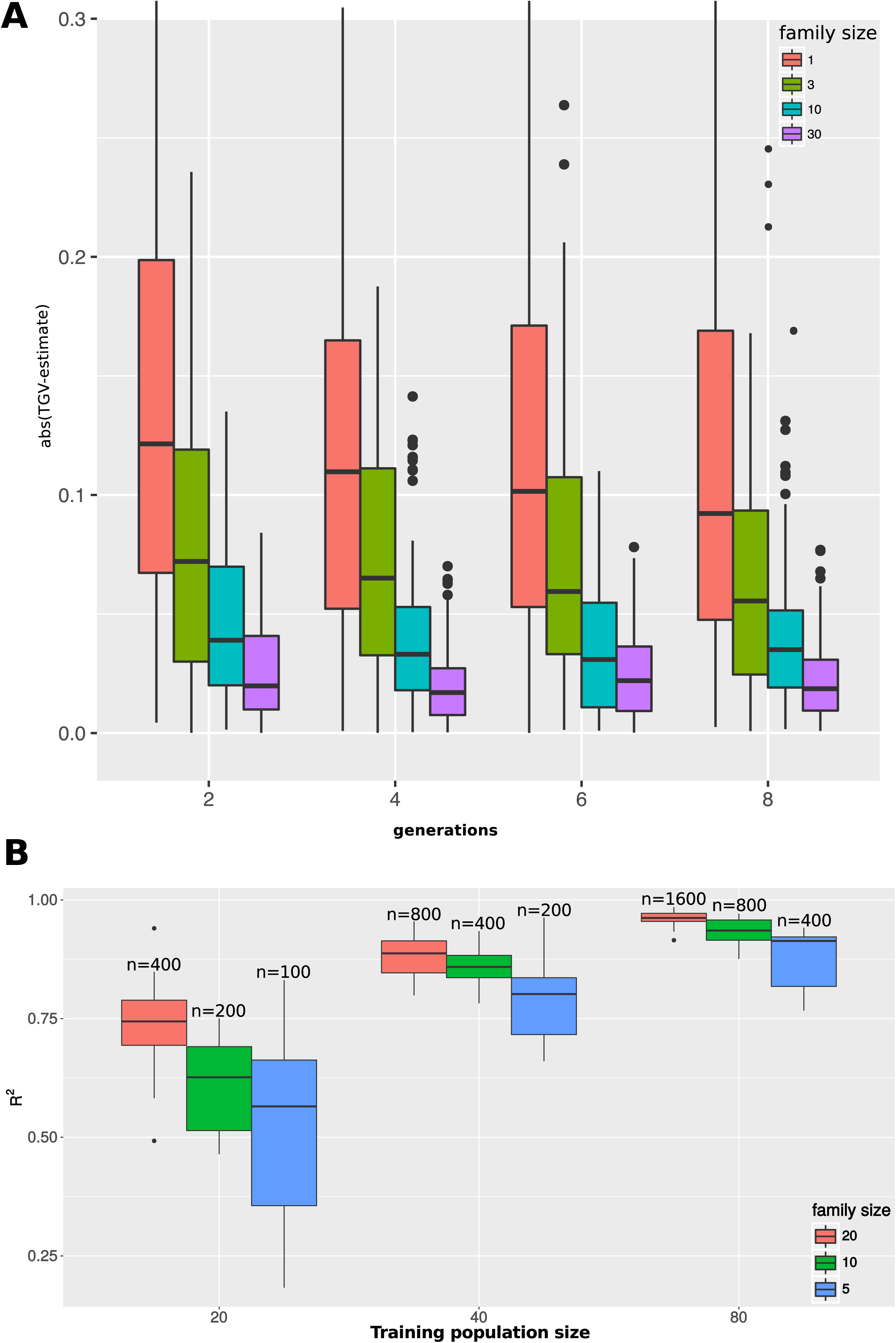
A) True breeding value estimation as a function of inbreeding and family replication. Boxplots depict the absolute difference between the TGV and the phenotypic estimate under a given set of parameters. The number of selfed generations are used to deivide groups along the X-axis and 4 different family sizes are shown for each generation level. B) Exploration of phenotypic accuracy versus training population size. Boxplots depict the prediction accuracy as an R2 value between the TGV of an individual and the genomic prediction of that individual. Models are based on the mean performance of families derived from individuals that are full-sibs of the test population.

Generally, replications are the measurement of one individual plant. Alternatively, some traits such as yield, are replicated as rows containing numerous plants. Thus, a few generations of advancement with bulking would be required in order to achieve the replication numbers of 10 and 30. If needed, *crossword* can easily simulate this possibility by advancing at the family level as opposed to the individual level (see Supplemental Figure 1).

Genomic selection (GS) is another area in which phenotypic accuracy is crucial. GS was developed within the cattle breeding community as a way of predicting complex traits solely from marker-based estimates of kinship to a phenotyped training set. Many crop breeding programs are now exploring best practices for the implementation of GS. In this context, one under-explored benefit of working in plants is the ability to phenotype genotypes over multiple years and locations to dramatically improve the estimate of the true genetic value. As with family homogenization and family size, optimal genomic selection logistics are additionally complicated by the need to determine the training population size that will result in the highest prediction accuracy given the available resources.

We simulated three different training population sizes and three different family sizes within each set (Figure 5B). “QTN_random” phenotyping method was used to sample 60 QTNs for each one of 10 iterations, and the QTN effect was assigned using gamma distribution. Populations were advanced to F5. We evaluated predication accuracy as the R^2^ between the TGV of an individual that has full-sibling relatedness to the training set and its phenotypic prediction based on the trained model.

Larger training set size is correlated with improved predication accuracy, although, at 400 replicates, sizes of 40 and 80 are roughly equivalent. Thus, it may be more appropriate for some species to have smaller training sets and larger family sizes depending on the species’ reproductive limitations. Even in large training sets, family size has a substantial impact, raising the median R^2^ by 10% with 5 extra replicates while also reducing the expected variation. Still, based on these results, as a “rule-of-thumb”, breeders should favor large training set over large family sizes.

### QTN polarity has a substantial impact on effect estimation of multigenic traits

In terms of simulating phenotypes, important varieties generally have some extreme trait that sets them apart. For example, one variety may be highly resistant to disease relative to another highly susceptible variety. Pre-existing platforms rarely support the ability to simulate this reality in an easy way. In *crossword*, QTN selection, aside from being supplied or random, can be biased towards one parent containing predominately positive or negative alleles. For mapping small number of QTN, this bias may be fairly irrelevant, but as the number of QTN increases, this scenario could substantially impact QTL detection and effect estimation.

We simulated four scenarios that highlight the consequences of asymmetrical effect polarity. In one set, 3 QTN and 15 QTN had randomly polarized effects (“QTN_random”) that are roughly evenly distributed between parents. Alternatively, in the second set (“highLow”), all effects are positive in one parent and negative in the other. For these simulations, we used a H^2^ of 0.7 and a population of 200 individuals advanced to F6 generations before phenotyping families of 5 individuals.

As expected, in both cases of 3 QTN, the polarity of the effects was irrelevant due to the fact that QTL are rarely in strong linkage with one another (Figure 6). As linkage becomes a factor with 15 QTN (~3 QTN per chromosome), effect estimates tend to vary more dramatically (Figure 6B). In the case of random polarity, fewer QTL should be detected because effects in the same linkage block will cancel each other out, creating the appearance of weaker, harder-to-detect QTL. This expectation is born out in simulation were the smallest number of QTL (67%) were discovered in the 15 QTN “random” mode and effects are generally underestimated (Figure 6B). Alternatively, when QTL have biased polarity in one parent (“highLow”), the QTL are more frequently detected (77%), and those missed are generally combined into a phantom peak. This combining is reflected in the fact that estimates are generally higher and some substantially exceed the actual effect size. Even in “random” mode, with 15 QTL, effect estimates can exceed the actual value by >15-fold, although this is a rare occurrence (1 out of 75 cases). Clearly, precise expectations for mapping an oligogenic trait can be very difficult to derive from first principles.

**Figure 6:**
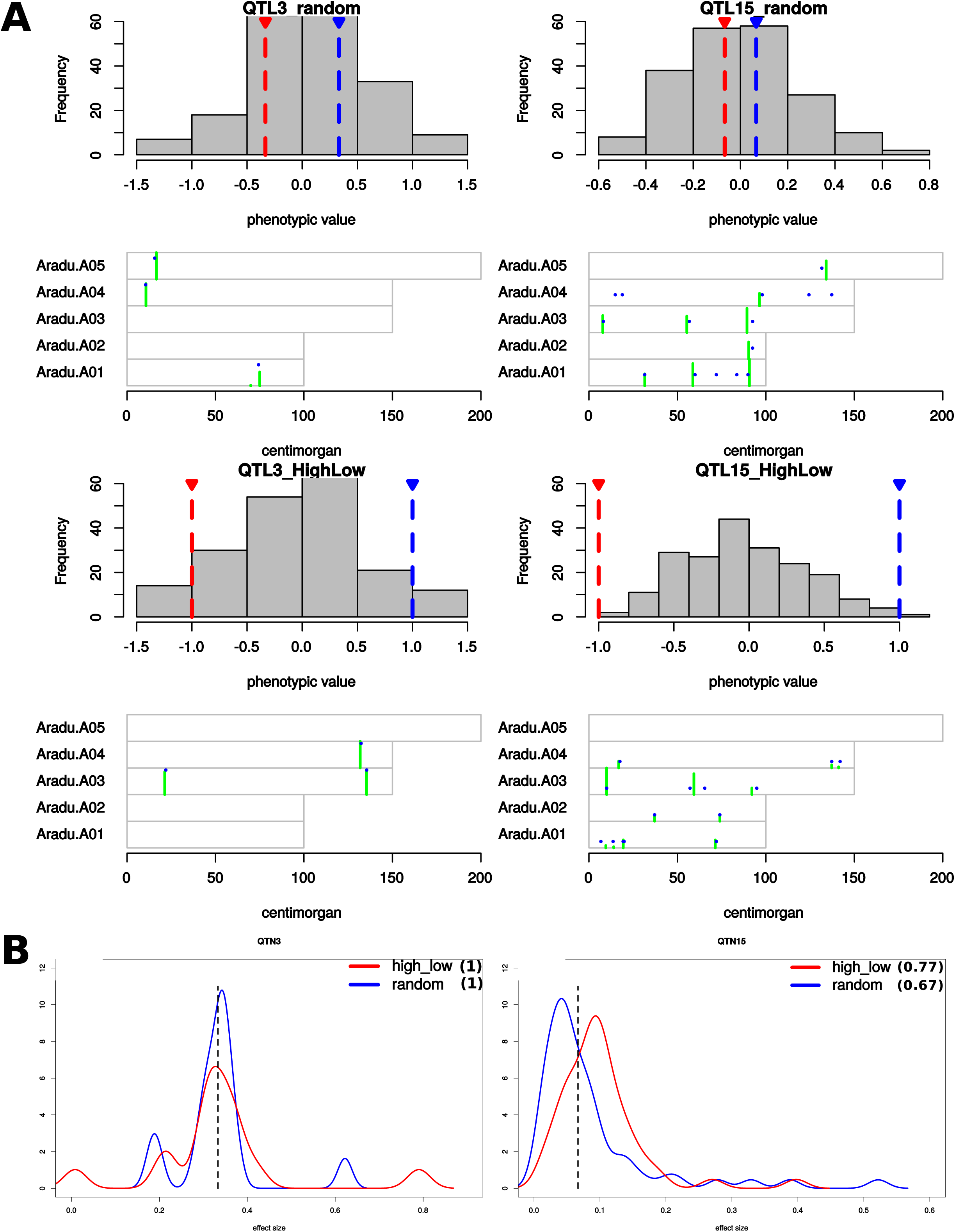
A) Impact of randomly polarized allelic effects versus parentally-biased. Each panel shows the phenotypic distribution of the mapping population and a chromosome plot of the simulated QTN position relative to the mapped position. In the distributions, high (red) and low (blue) parent values are indicated. In chromosome plots, blue dots indicate the simulated positions and effect sizes. (All the effects are equal.) Green bars indicate the estimated QTL position and the effect size. Chromosome maps are scaled in the y dimension to the largest effect size across all estimated effects across all chromosomes. B) Distributions of effect sizes across all iterations for highLow phenotyping methods with 3 and 15 QTN. The dotted vertical lines represent the actual effect for all QTN in the simulation. (3 QTN simulations have higher effect sizes than 15 QTN because maximum TGV in both cases is equal to 1.)

## Supplemental Figures

Supplemental Figure 1: crossword syntax overview with graphical examples.

Supplemental Figure 2: crossing-over loci for advanced generation of different population sizes.

